# Identification and Age-dependent Increase of Platelet Biased Human Hematopoietic Stem Cells

**DOI:** 10.1101/2022.01.14.475546

**Authors:** Merve Aksöz, Grigore-Aristide Gafencu, Bilyana Stoilova, Mario Buono, Yiran Meng, Niels Asger Jakobsen, Marlen Metzner, Sally-Ann Clark, Ryan Beveridge, Supat Thongjuea, Paresh Vyas, Claus Nerlov

**Author notes:** Equal contribution (Paresh Vyas, Claus Nerlov).

## Abstract

Hematopoietic stem cells (HSC) reconstitute multi-lineage human hematopoiesis after clinical bone marrow transplantation and are the cells-of-origin of hematological malignancies. Though HSC provide multi-lineage engraftment, individual murine HSCs are lineage-biased and contribute unequally to blood cell lineages. Now, by combining xenografting of molecularly barcoded adult human bone marrow (BM) HSCs and high-throughput single cell RNA sequencing we demonstrate that human individual BM HSCs are also functionally and transcriptionally lineage biased. Specifically, we identify platelet-biased and multi-lineage human HSCs. Quantitative comparison of transcriptomes from single HSCs from young, and aged, BM show that both the proportion of platelet-biased HSCs, and their level of transcriptional platelet priming, increases with age. Therefore, platelet-biased HSCs, as well as their increased prevalence and elevated transcriptional platelet priming during ageing, are conserved between human and murine hematopoiesis.

**One-Sentence Summary:** *In vivo* barcoding and single cell RNA sequencing identifies platelet-biased human bone marrow HSCs.

---

Hematopoietic stem cells (HSCs) sustain multi-lineage blood cell production throughout mammalian life. HSCs also reconstitute life-long blood cell production after hematopoietic cell transplantation (HCT), the most well-established and common form of clinical stem cell therapy (*1*). However, functional characterization of individual murine HSCs using single cell transplantation (*2-4*) and *in vivo* molecular barcoding (*5, 6*) has shown that in addition to HSCs with multi-lineage (ML) reconstitution capacity, a significant proportion of adult BM HSCs preferentially or exclusively generate a subset of hematopoietic lineages, and therefore are lineage-biased or fate-restricted (*1*). A key characteristic of murine fate-restricted HSCs is higher output of platelets compared to other lineages (*4*), and such HSCs are therefore frequently referred to as platelet-biased (P-) HSCs. Single cell transplantation has also demonstrated that ML-HSCs have higher overall cellular output compared to fate-restricted HSCs (*4*), a finding recapitulated in cellular barcoding experiments. These studies also identified transcriptional signatures associated with high vs. low cellular output and multi-lineage vs. platelet-biased lineage reconstitution (*7*). In the mouse, hematopoietic ageing is associated with significant expansion of P-HSCs (*8*). In contrast, the proportion of ML-HSCs, the only HSCs capable of lymphopoiesis, decreases with age, contributing to decreased adaptive immunity in aged individuals. Importantly, decreased lymphoid output has also been observed in aged human HSCs after xenografting into immune-deficient mice (*9*). Increased HSC transcriptional platelet priming may contribute directly to decreased lymphopoiesis during ageing, as inhibition of TGFβ signaling in aged HSCs leads to both decreased HSC platelet bias and increased lymphoid cell output (*10*). Finally, P-HSCs have been implicated as the cells-of-origin in a mouse model of essential thrombocythemia (ET) (*11*), and in human ET (*12*). In contrast to the mouse, characterization of the heterogeneity of human HSCs (hHSCs) is more limited. By xenografting, human cord blood (CB) with distinct long-term reconstitution capacity (*13, 14*) and repopulation patterns (*15, 16*) have been identified. These studies did not address hHSC contributions to the platelet and erythroid lineages, or include adult BM hHSCs. However, tracking of viral insertions in patients treated with HCT of gene corrected cells has identified single cell-derived reconstitution patterns consistent with the presence of fate-restricted hHSCs in adult BM (*17, 18*), and RNA sequencing revealed increased transcriptional platelet-lineage priming of aged compared to young BM hHSCs (*19*).

## Results

To identify putative lineage-biased hHSCs we performed high-throughput single cell RNA sequencing (RNAseq) on phenotypic bone marrow hHSCs (defined as LIN–CD34+CD38– CD90+CD45RA–; Fig S1) from 3 independent young adult donors (Table S1) using the 10X Chromium platform. A total of 13719 transcriptomes were assessed using area-under-the-curve (AUC) scores of transcriptional signatures associated with murine HSC lineage bias and output in individual hHSCs (*7*). This showed that multilineage- and high cellular output signatures were heterogeneously expressed and positively correlated at the single cell level, as were platelet-bias and low-cellular-output signatures, with individual HSC generating a continuum along both these axes (Fig. 1A). Furthermore, at the transcriptional level, platelet-bias was correlated to stemness and self-renewal signatures, whereas the multi-lineage signature was correlated to high-output and proliferation signatures (Fig. 1B), analogous to what has been observed in the mouse (*7*).

**Figure 1.**
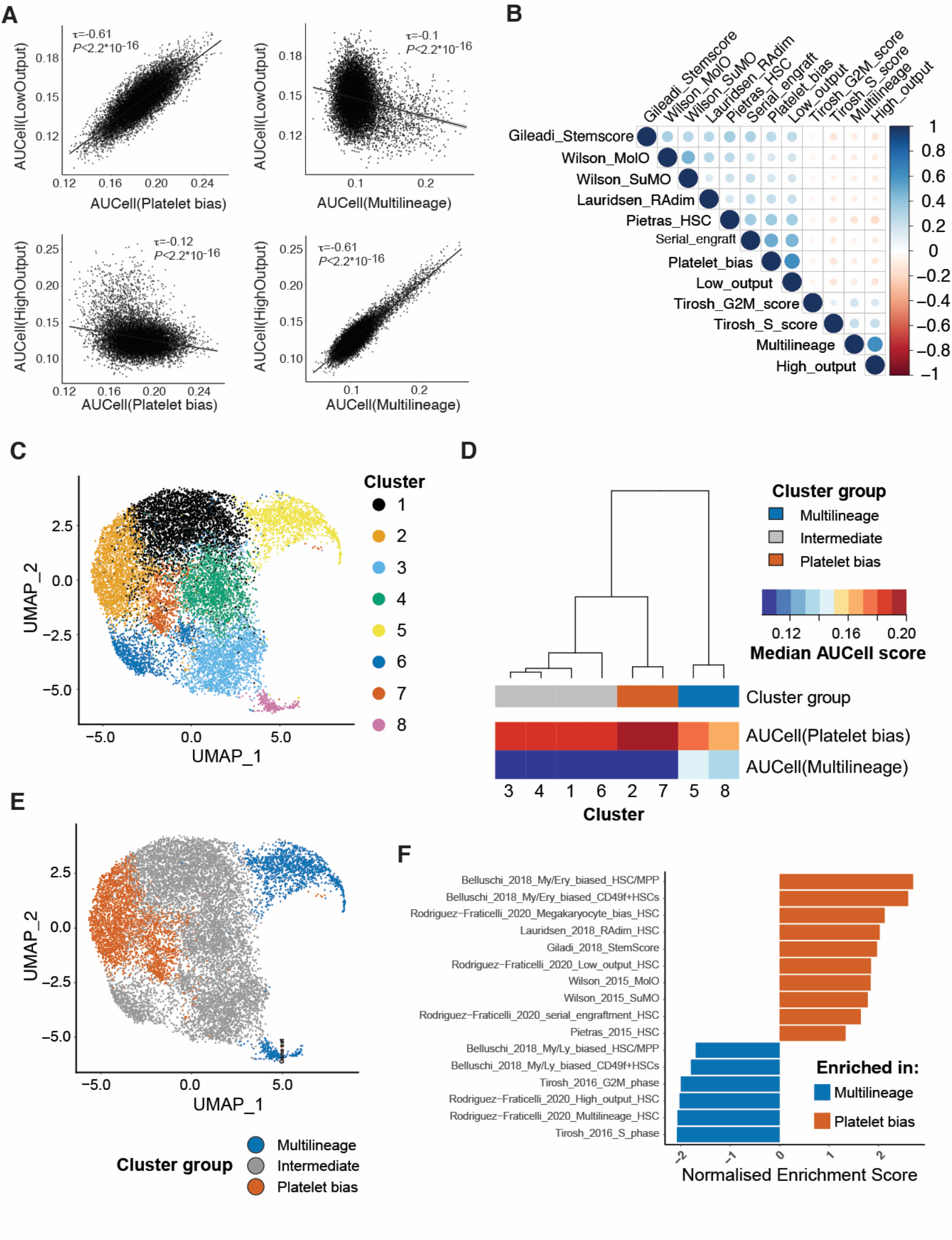
Computational identification of platelet biased HSCs in adult human bone marrow. **A)** Correlation between AUCell scores of the indicated transcriptional signatures defining HSC lineage bias and cellular output in transcriptomes of single human HSCs from adult bone marrow (N=13719, 3 independent donors). For each comparison the Kendal τ-b correlation coefficient and the associated P-value is shown. **B)** Kendal correlation of the indicated gene signatures from published datasets with in the single cell transcriptomes from (A). The scale shows the correlation coefficient. **C)** UMAP representation and clustering of the single cell transcriptomes from (A). **D)** Grouping of clusters from (C) by hierarchical clustering of using mean AUCell scores following unbiased clustering in cluster groups according to patterns of gene expression summarized by AUCell scores. **E)** Localization of the lineage-bias associated cluster groups in the UMAP representation from (C). **F)** GSEA normalized enrichment from comparison of the platelet-bias and multilineage cluster groups from (D) using the gene signatures from (B). Only gene signatures achieving a normalized enrichment score with FDR < 0.25 are shown.

To identify putative platelet-HSCs within the hHSC compartment we performed unsupervised clustering of single hHSCs (Fig. 1C), calculated the AUC scores for the lineage bias signatures for each cluster (Figure S2), and grouped these clusters based on their median platelet-bias and multi-lineage AUCell scores using hierarchical clustering (Fig. 1D,E). To determine if the putative P-hHSC populations had the expected molecular properties we compared their expression of lineage-bias and stemness/proliferation signatures to that of the putative ML-hHSC population. As expected, platelet-bias and low-output signatures were enriched in the putative P-hHSC population, as were signatures associated with stemness and quiescence; conversely, multi-lineage/lymphoid bias and proliferation signatures were depleted (Fig. 1F). The identification of putative P- and ML-hHSC populations also allowed the identification of genes differentially expressed between the two cell populations (Data S1).

To determine if hHSCs with functional lineage bias could be identified in adult BM, and whether the transcriptional profiles of computationally and functionally defined lineage-biased hHSCs were correlated, we combined barcoding and xenografting of BM hHSCs to simultaneously measure hHSC lineage output and characterize individual hHSC transcriptomes (Fig 2A). This involves transducing purified hHSCs (defined as above, Fig. S1) with a lentiviral barcode library (*20*) followed by xenografting into non-irradiated c-Kit–deficient immune-deficient NSG mice (NSGW41; (*21*)). The barcoded lentiviral vector expresses EGFP, allowing transduced cells to be identified and purified. Furthermore, as the barcode is transcribed as part of the proviral mRNA they can be identified by RNAseq of FACS-purified cells. Engraftment of hHSCs was verified by analysis of BM aspirates at 12- and 16-weeks post-transplant. We identified 7 mice, transplanted with HSCs from 3 different donors, with high levels of human BM engraftment (30-80%) and a significant proportion (10-40%) of transduced EGFP+ cells in the human BM fraction for subsequent analysis (Fig. S3).

**Figure 2.**
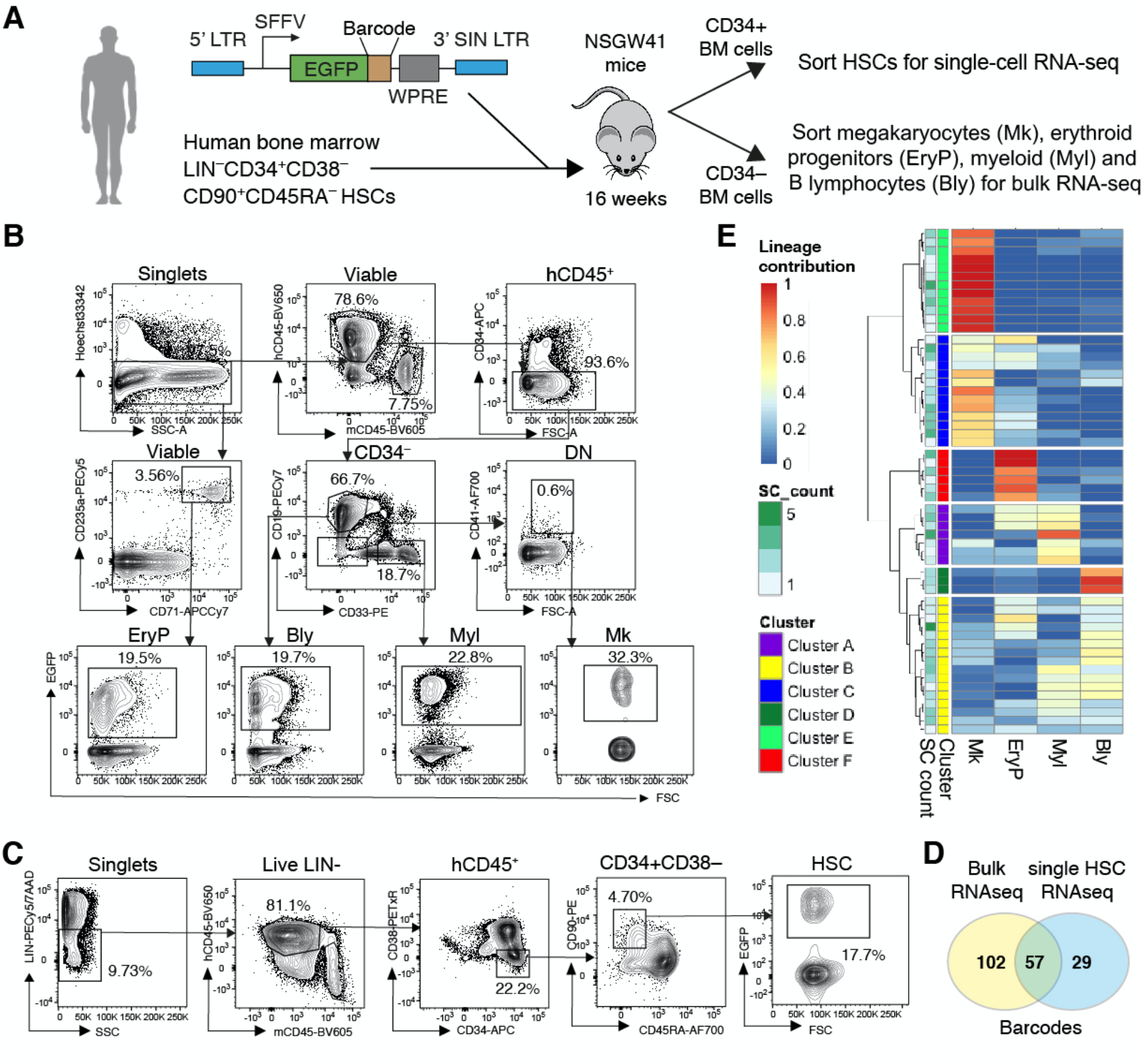
Lentiviral barcoding of adult human bone marrow HSCs. **A)** Experimental workflow and structure of the barcode lentiviral construct. 5’ LTR: 5’ long terminal repeat; SFFV: spleen focus forming virus; EGFP: enhanced green fluorescent protein; WPRE, Woodchuck hepatitis virus regulatory element; 3’ SIN LTR: 3’ self-inactivating LTR. The barcode is situated downstream of the EGFP and is expressed as proviral mRNA. **B)** Representative flow cytometry plots showing the gating strategy used to purify the cellular output of barcoded human HSCs from murine bone marrow. EryP: erythroid progenitor; Bly, B-lymphocytes; Myl: myeloid cells; Mk: megakaryocytes. Percentages shown are the proportion of the gated population. **C)** Representative flow cytometry plots showing the gating strategy used to purify barcoded human HSCs from murine bone marrow **D)** The overlap between cellular barcodes retrieved from bulk RNAseq of the lineage output isolated as in (B) and from single cell RNAseq of human HSCs retrieves as in (C). **E)** Heatmap showing hierarchical clustering of barcoded hHSC clones based on their lineage output. Each row represents an individual barcoded hHSC clone. SC count: shows the number of hHSCs within each barcoded clone Each column shows the contribution of the hHSC clone to different lineages (megakaryocyte, Mk; erythroid, EryP; myeloid, Myl; and B lymphocytes, Bly). The heat signature shows the normalized lineage contribution of each hHSC clone as fraction of total output. Based on lineage outputs, the hHSC clones fall into six clusters A-F, which are colour-coded as shown.

The engrafted mice were analyzed at the 16-week time point in order to focus on long-term repopulating hHSCs. BM cells were harvested and divided into CD34+ and CD34– fractions by magnetic cell sorting. From the CD34– fraction, EGFP+ cells were sorted for bulk RNAseq from the following populations: B-lymphoid cells (Bly; hCD45+CD34–CD33–CD19+), myeloid cells (Myl: hCD45+CD34–CD33+CD19–), erythroid progenitors (EryP; CD235+CD71+) and megakaryocytes (Mk; hCD45+CD34–CD33–CD19–CD41+SSChi) (Fig 2B). From the CD34+ BM fraction individual EGFP+ hHSCs were sorted (Fig. 2C) for single cell RNAseq. For both bulk and single cell RNAseq targeted amplification of the mRNA barcode was included in the RNAseq protocol. 251 barcodes were retrieved from the 4 lineage-restricted populations, of which 159 were represented by at least 20 barcode reads, the estimated number required for reliable detection of all lineages from a multi-lineage HSC (Fig. S4), and were retained in further analysis. From 472 single HSC transcriptomes 86 different barcodes were identified, each representing a clonal hHSC population. Of these 86 barcodes, 57 barcodes, representing 377 single hHSCs, were also detected in the bulk RNAseq (Fig 2D). These 57 hHSC clones were hierarchically clustered according to their lineage output, as determined by the distribution of the corresponding barcode in the bulk RNAseq of the 4 lineage-restricted populations. This identified 6 clusters with distinct output patterns, the 4 major clusters representing P-(cluster E), platelet-erythroid-(PE – cluster C), platelet-erythroid-myeloid-biased (PEM – cluster A) and ML (cluster B) patterns (Fig 2E). Similar output patterns were observed after transplantation of single murine HSCs (*4*), indicating conservation of HSC fate-restriction patterns between human and mouse.

Comparison of the transcriptomes of P- and ML-HSCs defined by barcoding identified genes differentially expressed between these HSC subtypes (Data S2). The genes up-regulated in P-hHSCs defined by barcoding (barcoded P-hHSC signature) showed significant overlap with those up-regulated in P-hHSCs identified by clustering (clustered P-hHSC signature) (P=4*10^−4^, χ^2^-test), and the signatures of P-hHSCs obtained by the two methods were significantly correlated at the single cell level across the hHSC population (Fig. 3B). Finally, comparison by GSEA of P- and ML-HSCs defined by barcoding and clustering, respectively, showed a similar enrichment pattern, with stemness and quiescence signatures enriched in P-hHSCs and proliferation signatures in ML-hHSCs (Fig. 3C). Therefore, the computationally and functionally identified hHSC populations were molecularly congruent.

**Figure 3.**
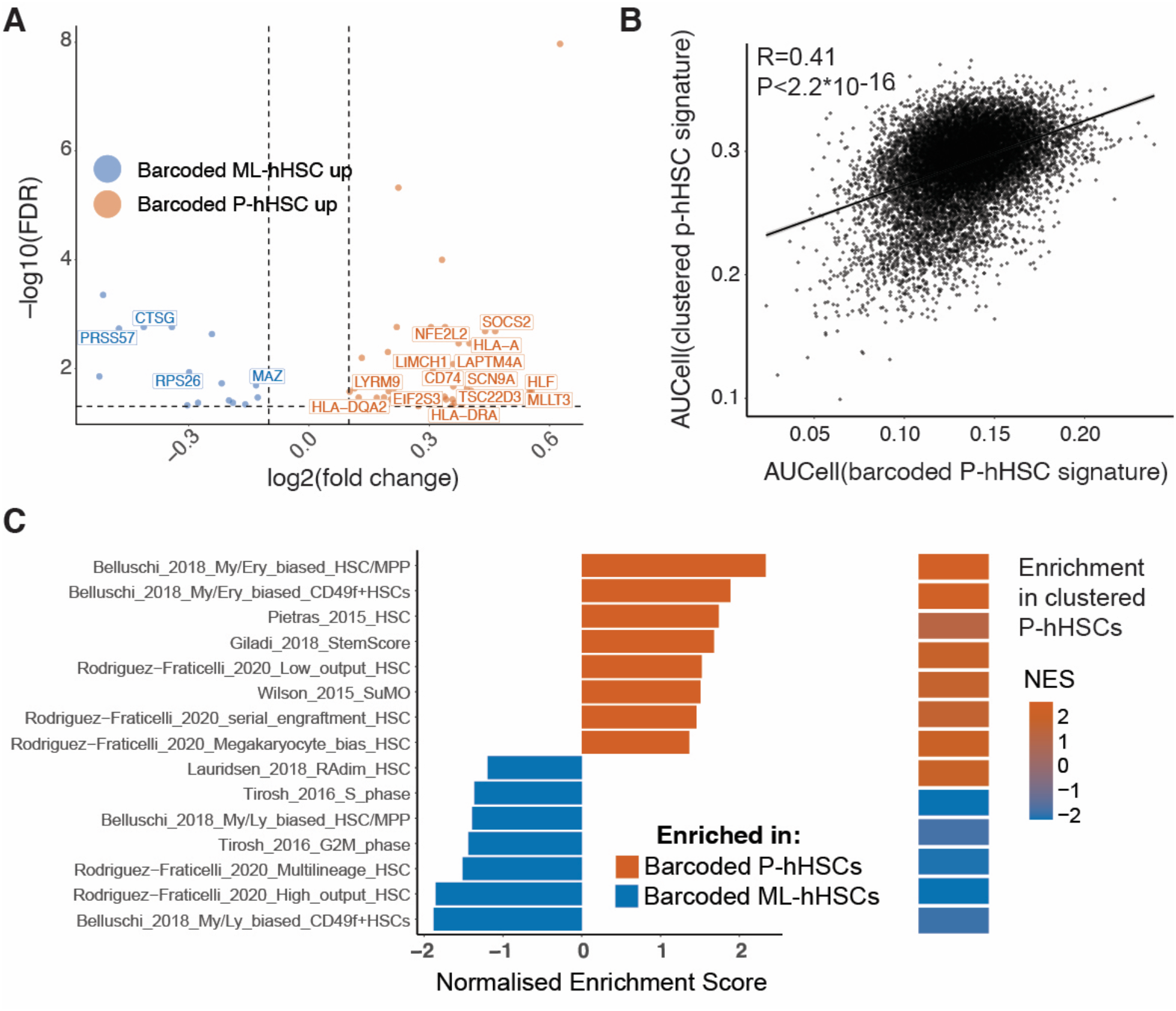
Comparison of platelet-biased HSCs identified by barcoding and computational clustering. **A)** Volcano plot showing genes differentially expressed between platelet biased and multilineage hHSC clones identified by barcoding (FDR<0.05, log2(fold change)>0.1). Gene IDs for genes also present in the P-hHSC and ML-hHSC signatures identified by clustering in Figure 1 are shown. **B)** Correlation between the AUCell scores of the P-hHSC signatures defined by barcoding (x-axis) and clustering (y-axis). The Pearson’s R correlation coefficient and associated P-value are shown. **C)** Comparison of P-hHSCs and ML-hHSCS defined by barcoding using GSEA of stemness and proliferation signatures from Figure 1B. Normalized enrichment scores are shown. Only gene signatures achieving an FDR<0.25 are included. The enrichment scores for the same analysis using the P-hHSC and ML-hHSC populations defined by clustering in Figure 1F is shown for comparison.

To address the effect of ageing on hHSC platelet-bias we next performed single cell RNAseq of aged hHSCs. A total of 10102 transcriptomes were generated from 3 aged donors (68-75 years old -Table S1). The expression of HSC output and lineage signatures showed the same correlation in young and aged hHSCs (Fig. S5A). However, aged hHSCs expressed higher levels of genes associated with platelet bias and lower levels of multi-lineage–associated genes compared to young hHSCs (Fig. 4A). To determine if this was due to an increase in the number of P-hHSCs the young and aged hHSC transcriptomes were co-clustered (Fig. 4B), and P- and ML clusters grouped by hierarchical clustering using the AUCell scores of platelet-bias and multi-lineage signatures as above (Fig. 4C,D, Fig. S5B). The enrichment pattern of stemness and proliferation signatures in old P- and ML-hHSC clusters (Fig. S6A) was conserved relative to young hHSCs (Fig. 1F), and similar analysis using canonical Hallmark gene signatures (MSigDB) also showed a high degree of conservation between young and old hHSC subtypes. In particular, metabolic (oxidative phosphorylation, fatty acid metabolism, glycolysis) and proliferation-associated signatures (including Myc targets, E2F targets, G2M checkpoint) were enriched in both young and old multi-lineage hHSCs, whereas inflammatory pathways (IL-6, TNFα, interferon, TGFβ signaling) and hypoxia-induced genes were enriched in both young and old P-hHSCs (Fig. S6B). Comparing the prevalence of aged and young hHSC in each cluster group showed that aged hHSCs were significantly enriched in P-hHSC clusters, and conversely depleted in ML-hHSC clusters (Fig. 4E). In addition, expression of the platelet-bias signature was increased in both platelet-biased and multi-lineage hHSCs during ageing (Fig. 4F). This indicates that ageing is associated with increased prevalence of platelet-biased hHSCs and a higher degree of platelet-lineage priming across all hHSCs, whereas differences in metabolic, proliferative and signaling pathways between hHSC subtypes were conserved during ageing.

**Figure 4.**
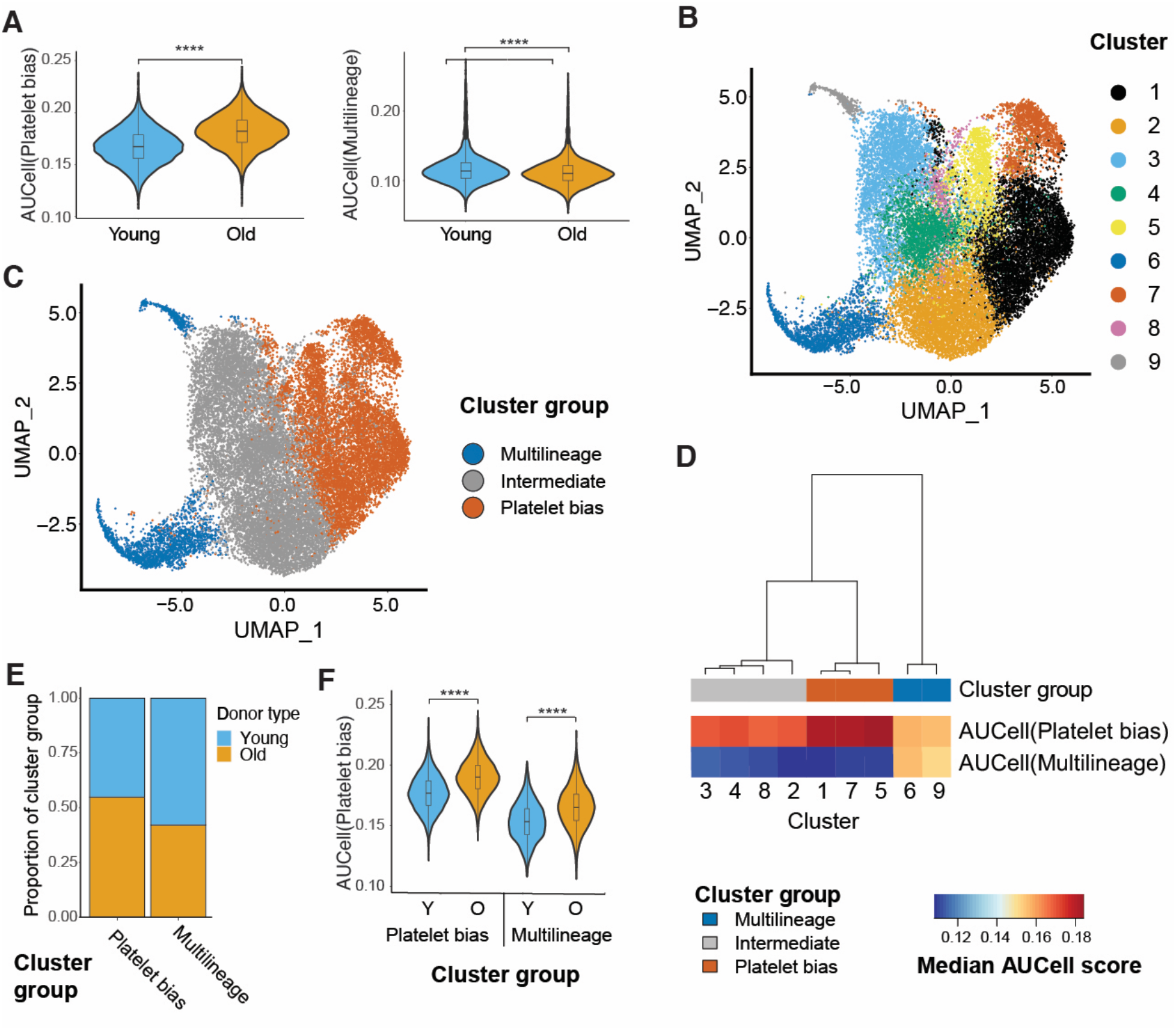
In silico identified platelet biased HSCs in human adult bone marrow exhibit an enrichment in old healthy donors. **A)** Distribution of AUCell scores for murine platelet bias (left panel) and multi-lineage HSC signatures (right panel) in hHSCs from young and old donors. ****: P < 0.0001 (Wilcoxon rank-sum test). **B)** UMAP representation and clustering of the single cell transcriptomes from young and old hHSCs. **C)** Grouping of clusters from (B) by hierarchical clustering of using mean AUCell scores following unbiased clustering in cluster groups according to patterns of gene expression summarised by AUCell scores. **D)** Localization of the platelet biased and multilineage cluster groups in the UMAP representation from (B). **E)** Proportion of young and old HSC in the platelet biased and multilineage cluster groups from (D). **F)** Distribution of AUCell scores murine platelet bias (left panel) and multi-lineage HSC signatures in young and old hHSCs from platelet biased and multilineage cluster groups in (D). ****: P < 0.0001 (Wilcoxon rank-sum test).

## Discussion

Studies using comprehensive readout of hematopoietic lineage outputs from individual murine bone marrow HSCs have consistently identified platelet- and platelet-myeloid biased HSC subtypes (*3, 4, 6*). However, parallel human studies have generally been performed using cord blood hHSCs, and not included measurement of the erythroid- and platelet lineages. By barcoding of adult hHSCs and comprehensive lineage readout we here find that hHSCs with similar fate-restrictions to those identified in the mouse (*4*), and P-hHSCs in particular, could be identified in human bone marrow. In addition, comparing the single cell transcriptomes of P- and ML-hHSCs, the differences in expression of stemness- and proliferation signatured were found to be conserved in hHSCs. P- and ML-hHSCs could also be identified using the previously defined murine HSC lineage-bias signatures (*7*), and P-hHSCs defined in this manner were transcriptionally similar to those identified by barcoding, with correlated expression of platelet-bias signatures and similar pathway enrichment. Therefore, both functionally platelet-biased hHSCs and their molecular programming are conserved between mice and humans.

Increased molecular and functional platelet bias is a key feature of aged mHSCs (*8, 19*), and this involves both increased prevalence of P-mHSCs and their increased platelet-lineage priming at the single cell level (*8*). We here find that both of these features of murine hematopoietic ageing are conserved in humans. Furthermore, using GSEA we identify a number of molecular pathways that are consistently enriched in young and old P-hHSCs. These include inflammatory signaling pathways and interferon and TGFβ signaling in particular. Increased interferon signaling has been observed in platelet-biased malignant hHSCs in essential thrombocythemia (ET), and our observations indicate that this is an inherent property of P-hHSCs that is conserved upon their transformation by *JAK2* mutation. In addition, P-hHSCs were enriched for hypoxia-associated gene expression compared to ML-hHSCs, which showed higher levels of genes involved in oxidative phosphorylation. P-mHSCs have previously been observed to occupy specific niches enriched in megakaryocytes (*22*), which produce Cxcl4 and TGFβ1 critical for maintenance HSC quiescence (*23*). The distinct inflammatory and metabolic signatures of P- and ML-hHSCs are consistent with P-hHSCs inhabiting similar niches within human bone marrow. Our results therefore indicate a high degree of conservation of HSC heterogeneity, the molecular properties of lineage biased HSCs, and the specialized micro-environments occupied by these between human and mouse.

## Acknowledgments

We thank the WIMM FACS facility for assistance with cell sorting, and the WIMM CBRG for computational support.

## Author contributions

M.A., B.S., M.B., Y.M., M.M. and S.A.C. performed the experiments. M.A., G.G., S.T. and C.N. analyzed the data, R.B. generated and validated the barcoded lentiviral library and barcode detection, N.A.J. collected bone marrow from aged individuals, C.N. and P.V. conceived the experiments, and C.N., P.V., M.A. and G.G. wrote the manuscript.

## Data and materials availability

Request for data, materials and code should be directed to C.N.

## Supplementary Materials

### Materials and Methods

#### Animals

The NSGW41 (NOD.Cg-Kit^W-41J^Tyr^+^Prkdc^scid^Il2rg^tm1Wjl^/ThomJ) mouse strain (*21*) was obtained from the Jackson Laboratory (Stock #026622) and bred in the animal facility at the University of Oxford (Biomedical Services). All experimental procedures, mouse breeding, and maintenance were performed according to UK Home Office regulations and with the ethical approval from the University of Oxford Medical Sciences Division Animal Welfare and Ethical Review Board.

#### Human bone marrow

Fresh BM from healthy young volunteers (21 to 34 years old males) were purchased from AllCells (Berkeley, CA) or Lonza (Lonza Bioscience, Basel, Switzerland). Bone marrow from older donors was obtained from orthoplastic surgery after informed consent. BM cells were filtered through 70 μm cell strainers (Corning), mononuclear cells (MNCs) were isolated Ficoll (GE Healthcare) density gradient centrifugation and CD34+ cell enrichment carried out according to the manufacturer’s protocol (Miltenyi Biotech).

#### Cell sorting

For sorting of human HSCs BM, CD34^+^ cells were stained with the following lineage markers (CD2, CD3, CD4, CD7, CD8, CD11b, CD14, CD19, CD20, CD56 and CD235a, all PE-Cy5 conjugated), CD34-APC, CD38-PE-Texas Red, CD90-PE, CD45RA-FITC, CD10-BV650, and CD123-PE-Cy7. 7-amino-actinomycin D (7AAD, Sigma Aldrich) viability dye was added to the samples prior to sorting to exclude non-viable cells. The clones and dilutions of all antibodies are listed in Table S2. Flow cytometry and cell sorting was performed using a BD FACSAria Fusion Cell Sorter with FACSDIVA software v.8.0.1 (Becton Dickinson and Company, Franklin Lakes, US-NJ). Data analysis was performed using FlowJo software (FlowJo LLC, v10.5.3).

#### Lentiviral barcoding and xenotransplantation

HSCs were transduced in SFEM serum-free media (Stem Cell Technologies) supplemented with human Flt3 ligand (100 ng/ml), KitL (100 ng/ml), Thpo (50 ng/ml), and IL-6 (50 ng/ml) (all from Peprotech). The peGZ2-linkerBC322 barcoding library (*20*) consisting of 725 barcodes was prepared as described (*24*). Purified HSCs were transduced at an MOI of 100-125 in StemSpan serum-free media supplemented with human cytokines Flt3 ligand, KitL, Thpo, and IL-6 as above for 18 hours at 37°C and 5% CO_2_, followed by intraosseous injection into 8-11 weeks old female NSGW41 murine recipients.

#### Purification of xenografted cells

BM cells were harvested from femurs, tibias, hips, sternum, and spine 16-weeks post-transplantations. Human CD34 magnetic separation then was performed to isolate CD34^+^ and CD34^−^ cell populations. CD34^+^ cells were stained as above with the addition of human CD45-BV650 and mouse CD45-BV605. Single cells were index-sorted into 96-well PCR plates (Thermo Scientific) containing 4μl of lysis buffer using a 85mm nozzle (*25*) for single cell RNAseq. CD34^−^ cells were stained with the following antibody panel: CD71-APC-Cy7, CD235a-PE-Cy5, CD19-PE-Cy7, CD33-PE, CD41-AF700, CD34-APC, human CD45-BV650 and mouse CD45 BV605. Hoechst 33342 was used as a viability marker. Barcoded human (hCD45^+^mCD45^−^) erythroid cells (EryP: CD235^+^CD71^+^EGFP^+^), B-cells (Bly: CD19^+^CD33^−^EGFP^+^), myeloid cells (Myl: CD19^−^ CD33^+^EGFP^+^), and megakaryocytes (Mk: CD41^+^CD19^−^CD33^−^EGFP^+^) were sorted directly into PCR tubes containing 4μl of lysis buffer (50 or 100 cells/biological replicate) for bulk RNAseq using a 130μm nozzle.

#### Single cell and bulk RNA sequencing library preparation

Single-cell and bulk libraries were generated using Smart-seq+ (single cell RNAseq) (*26*) and targeted Smart-seq2 (bulk RNAseq) (*27*), respectively. Custom primers for barcoding amplification were included during the initial cDNA preparation and pre-amplification steps. The sequences of the primers used in single-cell and bulk RNA-seq is provided in Table S3. 22 cycles were performed for PCR amplification. After pre-amplification, cDNA libraries were purified using Ampure XP (Beckman Coulter) magnetic beads according to the manufacturer’s instructions, in a ratio of 1.2 to 1 with cDNA. Following tagmentation and 12-cycles of index PCR using Nextera adapters (Illumina), libraries were bead-cleaned up using a 1:1 ratio of Ampure XP beads (Beckman Coulter). All cDNA and indexed libraries were quality checked for their fragment sizes using Agilent high-sensitivity chips on an Agilent 2100 Bioanalyzer (Agilent Technologies) or an Agilent fragment analyzer high-sensitivity kit with an Agilent fragment analyzer (Agilent Technologies). Library concentrations were assessed using a Qubit dsDNA HS kit (Invitrogen) and a Qubit fluorometer (Invitrogen). Dual-indexed libraries were pooled to a final concentration of 2 nM and sequenced on an Illumina NextSeq500 instrument (75-bp single-end reads) using a NextSeq 500/550 High Output kit v2.5 (Illumina).

#### Single cell 10X Chromium library preparation

HSCs were purified from young and aged human BM (N=3 for both ages; Table S1). Libraries were prepared using the Chromium Single Cell 3’ Reagent Kits v2 (10X Genomics, USA). cDNA concentrations were assessed using a Qubit dsDNA HS kit (Invitrogen) and a Qubit fluorometer (Invitrogen). Fragment sizes were checked using an Agilent high-sensitivity chip (Agilent Technologies) on an Agilent 2100 Bioanalyzer (Agilent Technologies) according to the manufacturer’s instructions. Libraries were sequenced on a NovaSeq6000 S4 flow cell (NovoGene).

#### Single cell and bulk RNAseq analysis

Nextera adapter sequences and low quality reads were removed from demultiplexed FASTQ files using Trim Galore (v0.5.0) (*28*). Reads were aligned to the hg38 human reference genome using STAR (v2.6.1d) (*29*). Unique read counts were quantified using featureCounts (subread/v1.6.2) (*30*). Quality control (QC) metrics were generated using MultiQC (v0.9) (*31*).

#### Barcode detection and analysis

The merged FastQ files were used for the barcode detection in single-cells and bulk samples. The barcodes in HSCs and their lineage outputs were identified using a custom R script to quantify the barcode count number. All reads were searched for sequence matching the following barcode pattern: NNNACNNNGTNNNCGNNNTANNNCANNNTGNNN. The biological replicates of Bly, Myl, Mk, and EryP lineages were aggregated and the total barcode reads for each barcode and each lineage was calculated. To estimate the number of barcode reads required for reliable detection of all lineages generated by an hHSC clone a random draw was simulated from a population of 10^6^ equally distributed across N subpopulation (N being the number of lineage generated by the clone, ranging from 2-4). 10^5^ simulated draws were carried out, and the frequency of detection of 1 to N lineages calculated. The cutoff was set at a 99% chance of detecting all 4 lineage generated by a multi-lineage hHSC (20 reads – see Figure S6). Barcodes with less than 20 total reads were therefore filtered out. The barcode frequencies were calculated by dividing the barcode sum values by the total aligned reads of their respective lineage. This normalizes the barcode counts to the sequencing depth. Next, for a reliable comparison of the lineages, the barcode detection level was normalized, as barcodes make up different fractions of the transcriptome of different lineages. For this reason, scaling factors were generated using the multilineage clones (clones with barcode reads for all lineages, where the average contribution of the four lineages was assumed to be even, as observed in mouse multilineage clones) as a reference and used to normalize the clone barcode counts across lineages. To classify HSC clones by their fate, the normalized barcode reads were divided by their sum for each clone, and the resulting scaled values were used for hierarchical clustering of HSC clones using the R dist function to calculate Euclidean distance and the R hclust function to perform wardD2 agglomeration. Clusters were identified by cutting the dendrogram to six clusters using the R cutree function.

#### Chromium 10X scRNA-seq data processing and quality control

The raw fastQ files were aligned against the *H. sapiens* GRCh38 (Ensembl 93) reference genome (10X Cell Ranger reference GRCh38 v3.1.0) and quantified recommended using the Cell Ranger pipeline (v3.1.0) at default parameters. The *count* functionality of the Cell Ranger pipeline was utilised to generate the filtered cell x gene raw UMI family counts for the data from each sample individually and further processed using Seurat (v4.0.1) (*32*). All data has been subjected to the same QC filtering: cells with <200 genes detected and >10% of their UMIs mapping to mitochondrial genes were not considered for further analyses and data from genes that were not detected in at least 3 cells was discarded. Data that passed QC was normalised using the Seurat’s *scTransform* command, which implements regularized negative binomial regression for normalization and variance stabilization. In order to mitigate the batch generated during the library preparation procedures, each sample being considered a batch, after normalization, 3000 integration anchors across samples were defined using the Seurat’s *FindIntegrationAnchors* function. Once the integration anchors were defined, data was batch corrected using the Seurat *IntegrateData* function, setting normalization.method = “SCT”.

#### Dimensionality reduction and clustering

Following the batch correction of the data, principal component analysis (PCA) was conducted using Seurat’s *RunPCA* function the data from the 3000 integration anchor genes previously identified for the batch correction procedure. Using an elbow plot describing a ranking of principle components based on the percentage of variance explained by each one, the first 12 PCs were used as the input for the Uniform manifold approximation and projection (UMAP) visualisation performed using Seurat’s *RunUMAP* function of the dataset representing the young HSC data and the first 15 PCs for the analysis of the dataset integrating the HSC data from the young and old donors. Clustering of the data was carried out by first constructing a shared nearest-neighbour (SNN) graph, with edges drawn between the cells with similar gene expression patterns using an algorithm implemented in Seurat’s *FindNeighbors* function at default parameters and utilising as input the gene expression data summarised by the first 12 PCs for the young HSC dataset and the first 15 PCs for the combined young & old HSC dataset analysis. Using the previously computed SNN graph, the cells were clustered using the Louvain algorithm’s implementation in Seurat’s *FindClusters* function used at default parameters. The resolution parameter in the *FindClusters* function applied in the combined young & old HSC dataset analysis that yielded the most robust clustering (resolution = 0.5) was determined following a previously described methodology (*33*). Clusters were assigned into clusters groups, reported as P-hHSCs or ML-hHSCs, based on the hierarchical clustering of the Euclidean distance computed for each cluster’s median platelet bias & multilineage AUCell scores using the ward.D2 agglomeration method as implemented in stats R package (v4.0.5).

#### Differential gene expression analysis

Differentially expressed genes between distinct cluster groups or HSC clone types were identified using Seurat’s *FindMarkers* function using the following parameters changed from the default ones: logfc.threshold = 0, test.use = “MAST”, min.pct = 0, assay = “RNA”. The p-values of the for each gene entry in the output of the Seurat’s *FindMarkers* function were corrected for multiple comparisons using the Benjamini & Hochberg correction as implemented in the *p*.*adjust* function of the stats R package (v4.0.5). Genes were considered differentially expressed between clusters or HSC clone types if they had a corrected p-value < 0.05.

#### Gene expression score computation and assessment

Gene expression scores for the gene sets of interest were computed using AUCell module of the SCENIC suite for scRNA-seq data analysis (*34*). The gene symbols for the gene sets used for computing AUCell scores (*7, 35-39*) were converted into their human counterparts when possible by utilizing the *getLDS* function as implemented in biomaRt (v2.46.3) (*40*) and utilising the *H. sapiens* (GRCh38.p13) and *M. musculus* (GRCm39) Ensembl gene symbol annotation for the gene symbol conversion. The gene lists used to compute the AUCell scores in the young donor HSC dataset were obtained by intersecting the aforementioned gene lists after the gene symbol conversion procedure (Data S3) with the list of highly variable genes determined using Seurat’s *scTransform* command on the data from the young donor HSCs, with the exception of the gene lists pertaining cell cycle phase progression (*37*) which were used without this filtering step. The gene lists used to compute the AUCell scores in the combined young & old donor HSC dataset were obtained by intersecting the gene lists generated after the gene symbol conversion procedure (Data S3) with the list of the integration anchor genes used for batch correction across samples in the combined young & old HSC dataset, with the exception of the gene lists pertaining cell cycle phase progression (*37*) where the gene lists were used without this filtering step. The genes upregulate in the platelet bias cluster group HSCs identified in the young donor HSC dataset that were expressed in at least 1% of the platelet bias cluster group HSCs or the multilineage cluster group HSC and had an FDR < 0.05 (n=774, see Data S1) were used to compute the clustering platelet bias AUCell score and the genes upregulate in the barcode platelet bias HSCs with an FDR < 0.05 (n=62, see Data S2) were used to compute the barcode platelet bias AUCell score. Following the processing of the gene sets, for each cell in the dataset the genes detected in each cell were ranked according to their expression from the ones with the highest expression to the one with the lowest one using the AUCell’s *AUCell_buildRankings* function. Next, for each cell in the dataset in order to determine if the gene sets of interest are enriched in the top ranked genes according to their expression, the AUCell’s *AUCell_calcAUC* function was used at default parameters. The differences in gene expression score distributions across the clusters determined in the datasets as well as the type of donors were evaluated using the Wilcoxon rank-sum test, as implemented in the *pairwise*.*wilcox*.*test* and *wilcox*.*test* function in the stats R package (v4.0.5). When comparing gene expression scores between 3 or more clusters, the significance of the differences in distributions observed was quantified by using the Wilcoxon rank-sum test in a pairwise fashion and the p-value of the test was corrected for multiple comparisons using the Benjamini & Hochberg correction as implemented in the *p*.*adjust* function of the stats R package (v4.0.5). The Pearson’s R or Kendall’s τ correlation coefficients were computed using the *cor*.*test* function of the stats R package (v4.0.5) in order to quantify the strength and direction of association between 2 types of AUCell scores; the choice of one type of coefficient over the other was predicated on the normality of the distribution of the AUCell scores investigated in. this study.

#### Differential cell type abundance analysis

In order to analyse if the size of a cluster group is dependent on the HSC donor type in terms of age group, the Pearson’s χ^2^ test as implemented in the R (v4.0.5) *chisq*.*test* function was used.

#### Gene set enrichment analysis

Gene set enrichment analysis (GSEA) was conducted starting from the output of Seurat’s *FindMarkers* function and ranking the full list of gene average fold changes per HSC cluster group or clone obtained by comparing the normalised UMI/read counts from different cluster groups or clones by their magnitude and sign. The enrichment of the gene lists of interest, including the murine gene lists whose gene symbols have been converted into human ones for the computation of AUCell scores (*7, 35, 37-39, 41-43*) (Data S3) was evaluated in the ranked gene lists generated following the differential gene expression analysis by using the *fgsea* wrapper function of the fgsea R package (v1.18.0) (*44*) with the following parameters: minSize=5, maxSize=1500, eps = 0, nPermSimple = 100000. Assessing if the overlap between gene signatures was statistically significant in terms of its magnitude was evaluated using the *newGOM* function of the GeneOverlap R package (v1.28.0) (*45*) that implements the Fisher’s exact test to perform the hypothesis testing and the p-values for each comparison were corrected using the Benjamini & Hochberg method.

## Supplementary Figures

**Figure S1.**
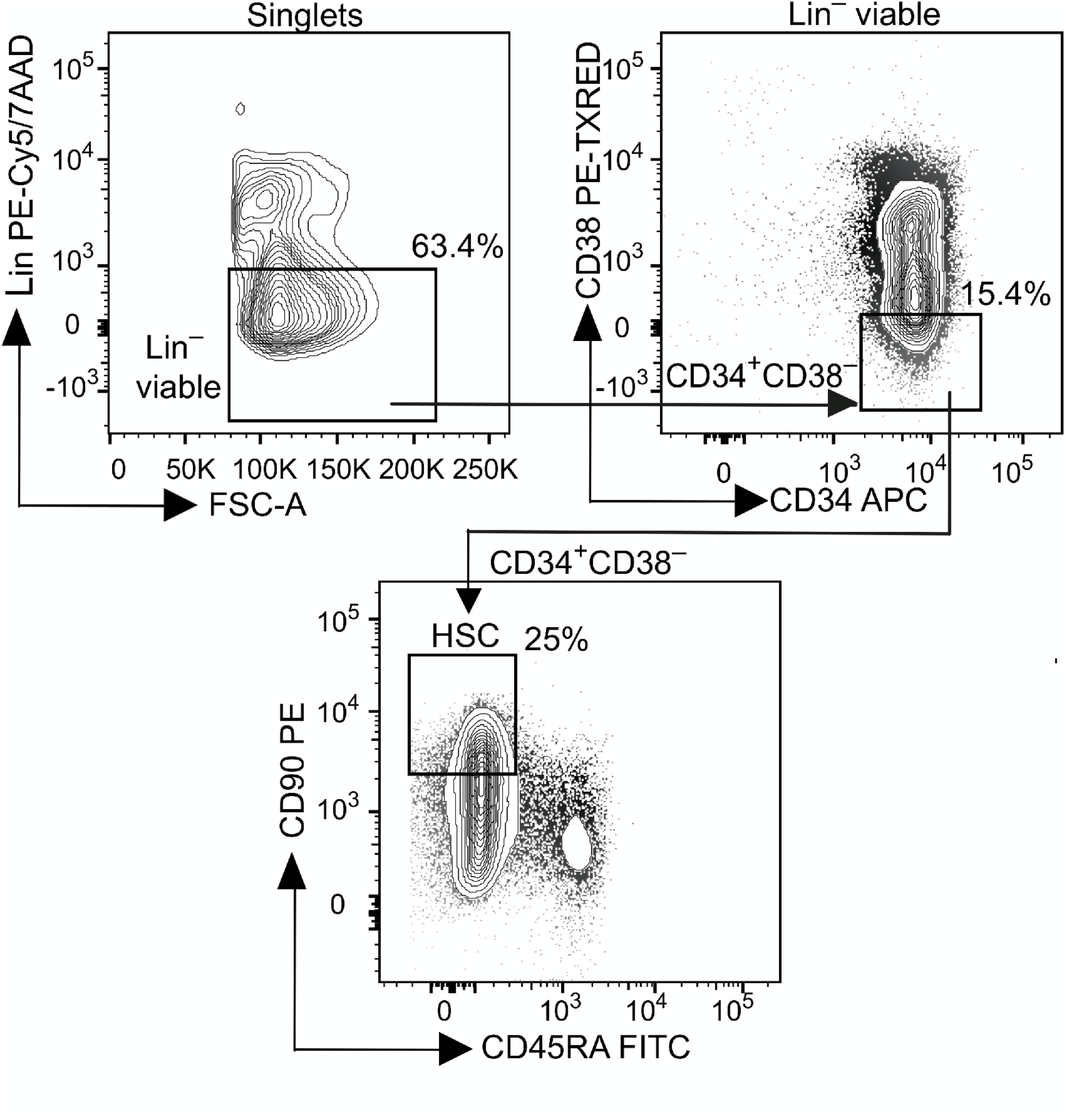
Gating strategy for bone marrow hHSC purification. Human bone marrow hHSCs were FACS purified using the sorting strategy outlined. The gated populations as percentage of the parental gate are show.

**Figure S2.**
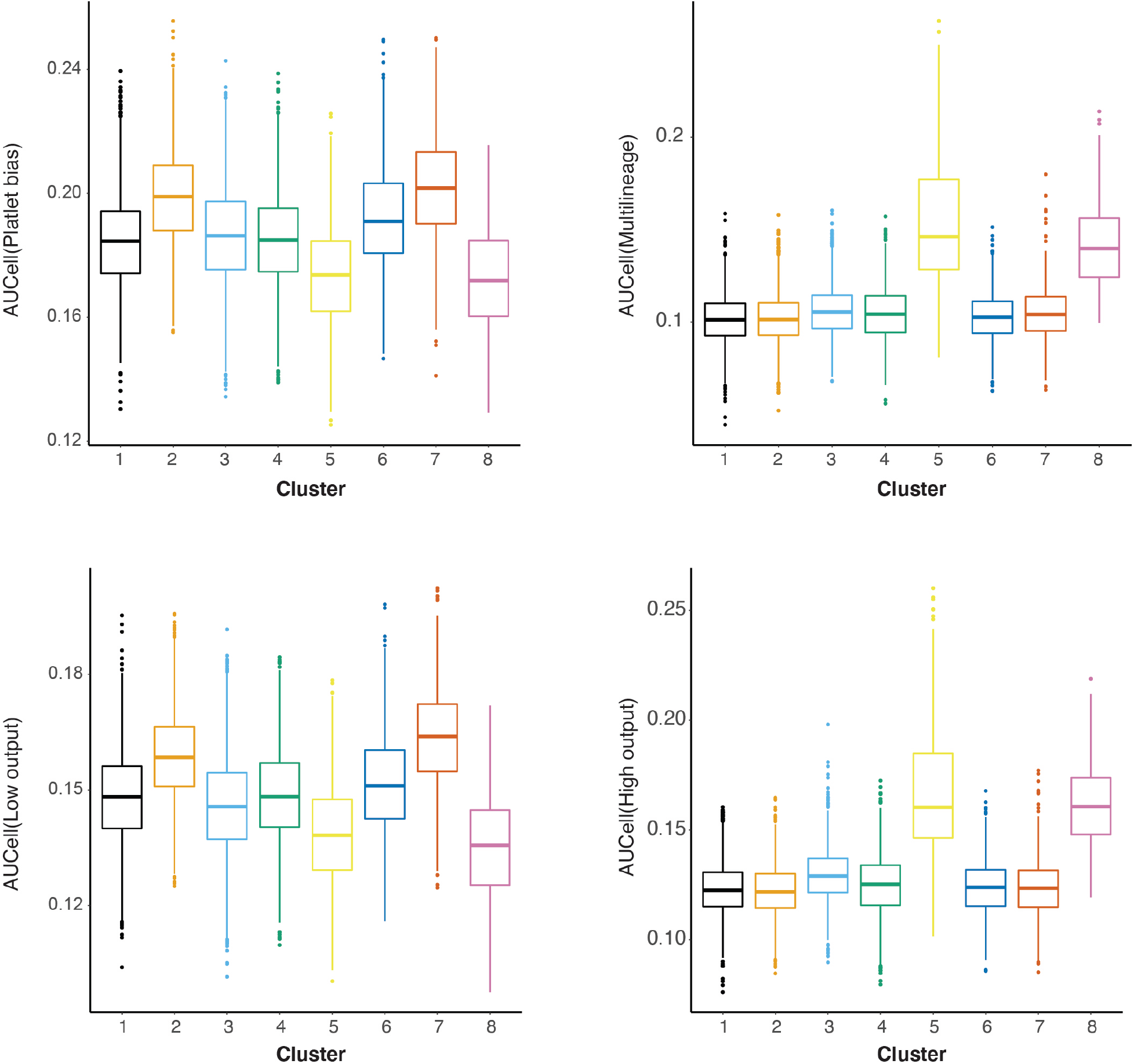
AUCell score distribution for the indicated gene signatures in the hHSC clusters defined in Figure 1C.

**Figure S3.**
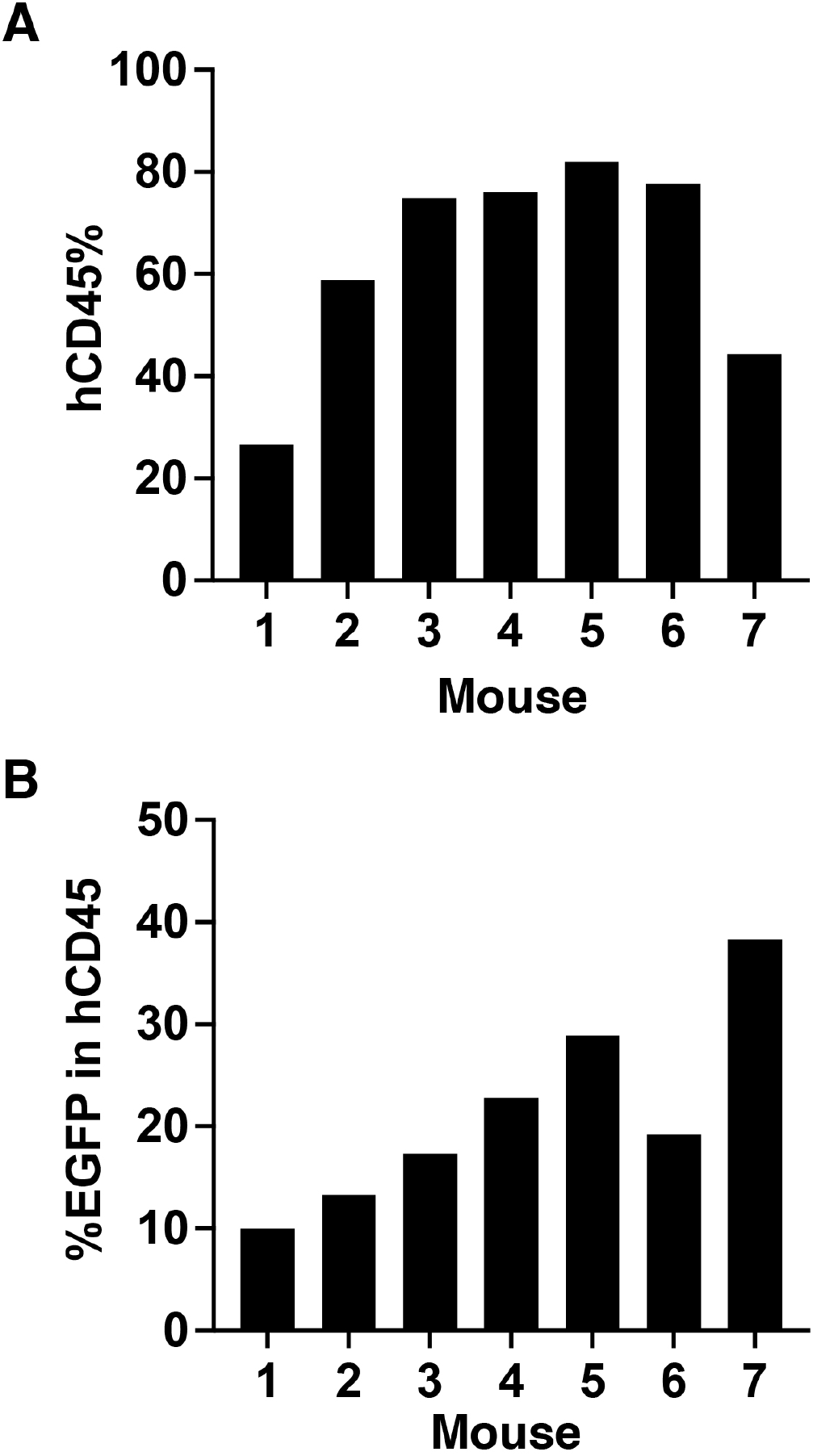
Human engraftment and transduction efficiency in xenografted mice. **A)** Human chimerism within the murine bone marrow leukocyte population from mice used for barcode analysis. The bars show the percentage of hCD45+ cells within the viable singlet gate as defined in Figure 2B. **B)** Bar graph showing the percentage of EGFP+ cells within the hCD45+ population, as defined in (A).

**Figure S4.**
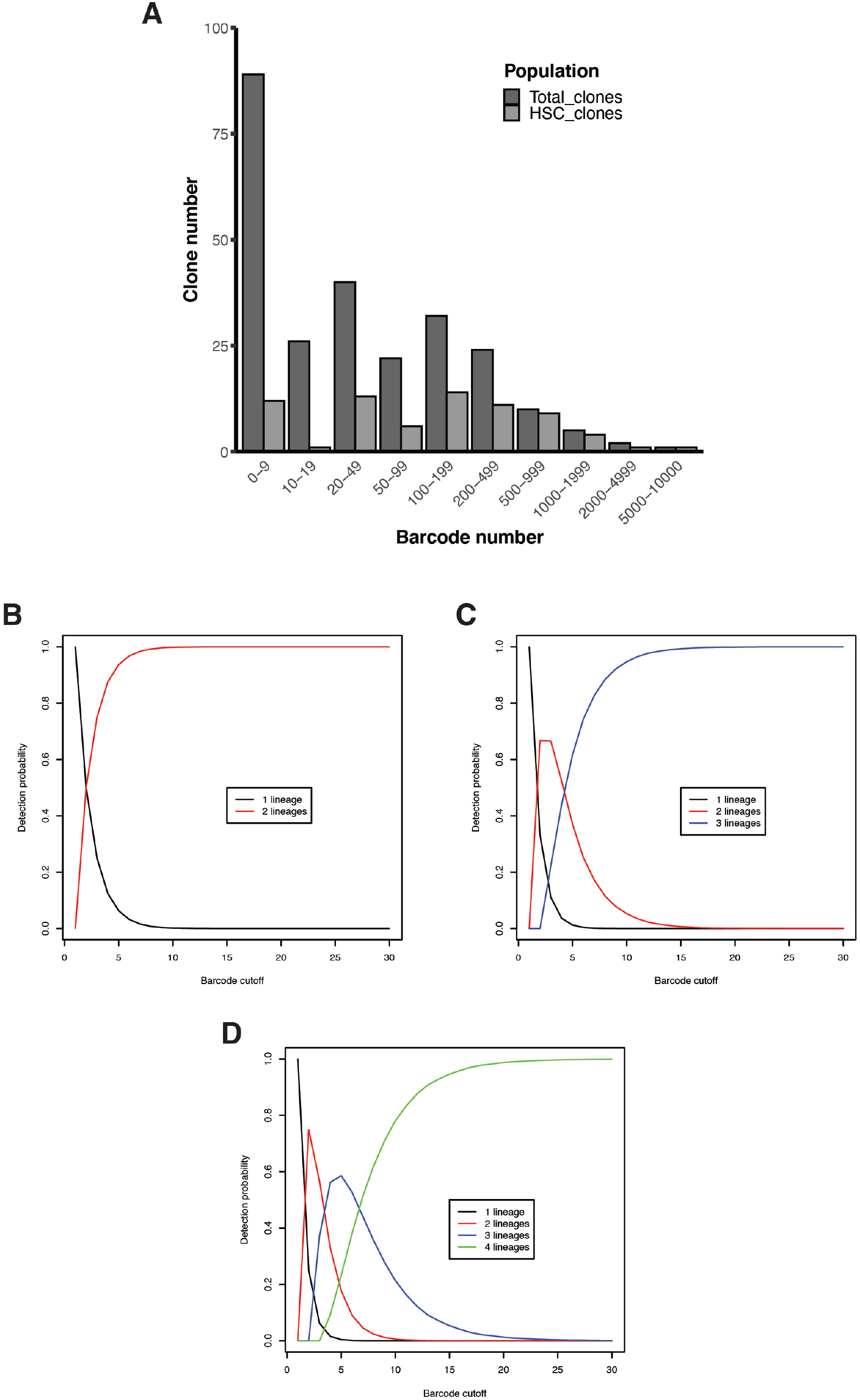
Distribution of barcode reads in HSC clones. **A)** Bar graph showing the number of barcodes detected with a number of reads within the indicated intervals by the bulk RNAseq (total clones), as well as the number of these barcodes that were detected in the single cell RNAseq of purified HSCs. **B)** Modeling of the probability of detecting one or two lineages from an HSC clone with two lineage outputs as a function of the number of detected barcode reads, using random sampling and assuming equal probability of barcodes being found in each lineage. **C)** Modeling of the probability of detecting one, two or three lineages from an HSC clone with three lineage outputs as a function of the number of detected barcode reads, calculated as in (B). **D)** Modeling of the probability of detecting one, two, three or four lineages from an HSC clone with four lineage outputs as a function of the number of detected barcode reads, calculated as in (B).

**Figure S5.**
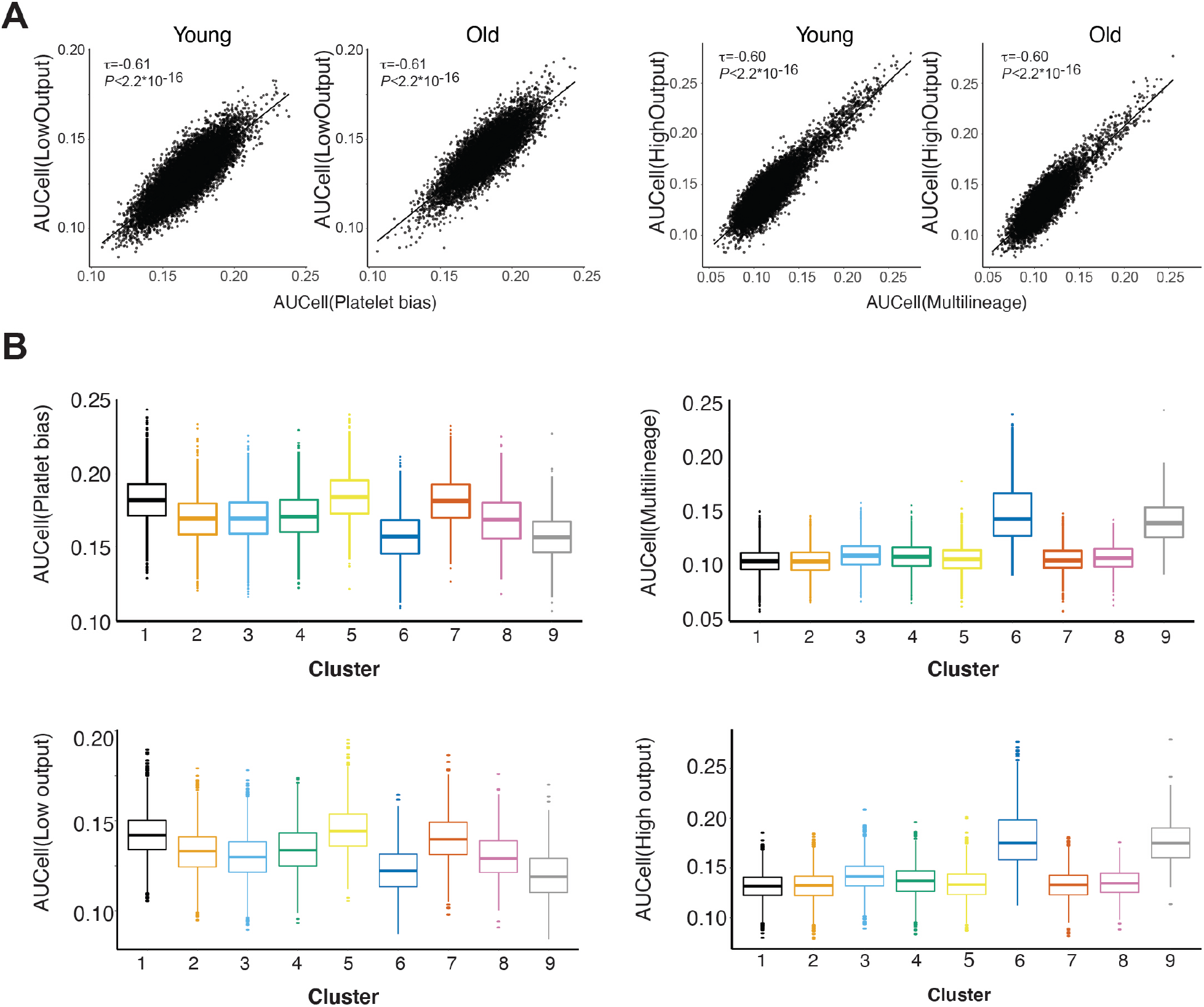
Analysis of combined young and old hHSC datasets. **A)** Correlation between AUCell scores of the indicated transcriptional signatures defining HSC lineage bias and cellular output in transcriptomes of single human HSCs from young (left panel) and old (right panel) adult bone marrow (N=13884, N=10301, respectively, 3 independent donors/condition). For each comparison the Kendal τ-b correlation coefficient and the associated P-value is shown. **B)** AUCell score distribution for the indicated gene signatures in the hHSC clusters defined in Figure 4B.

**Figure S6.**
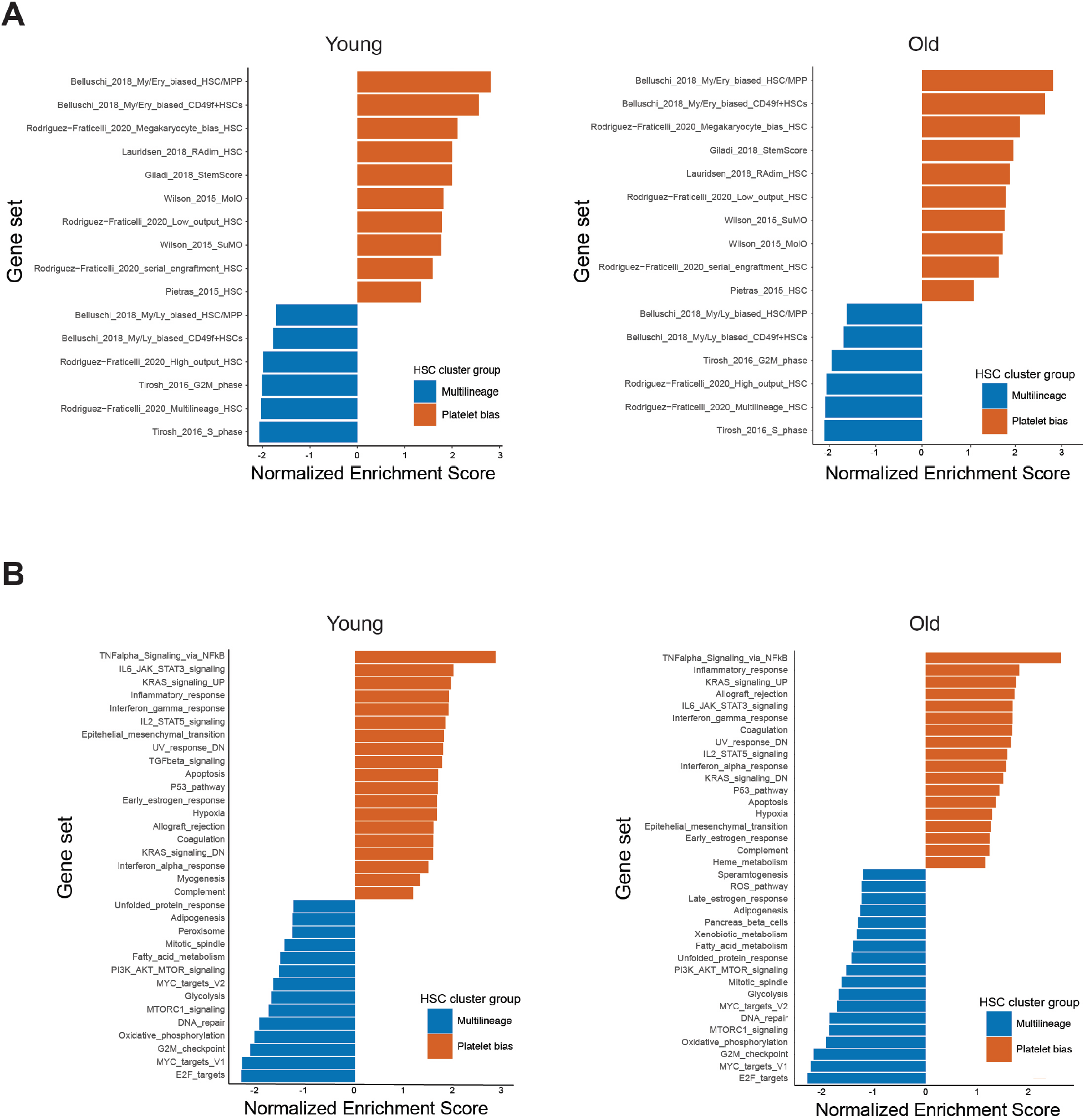
Comparison of young and old lineage-biased hHSCs by GSEA. **A)** hHSC subtypes defined by clustering in Figure 4B were divided into young and old subsets, and platelet-biased and multilineage hHSCs were compared for both ages using GSEA and stemness/proliferation signatures from Figure 1B. Gene signatures achieving a normalized enrichment score with an FDR < 0.25 are shown. **B)** Analysis as in (A) using the MSigDB Hallmark pathway gene set.

## Supplementary Tables

**Table S1.** Donor used for 10X Chromium sequencing.

| **Donors** | **Age** | **Gender** | **Number of cells purified** | **Total recovered cells** | **Mean reads per cell** | **Cells post QC** |
| --- | --- | --- | --- | --- | --- | --- |
| 1 | 21 | Male | 12,000 | 4,327 | 172,155 | 4,299 |
| 2 | 24 | Male | 12,000 | 3,909 | 136,597 | 3,888 |
| 3 | 22 | Male | 12,000 | 5,648 | 94,207 | 5,532 |
| 4 | 68 | Male | 13,130 | 4,684 | 107,314 | 4,579 |
| 5 | 71 | Male | 6,658 | 2,638 | 97,334 | 2,574 |
| 6 | 75 | Male | 7,078 | 2,989 | 87,326 | 2,949 |

**Table S2.**
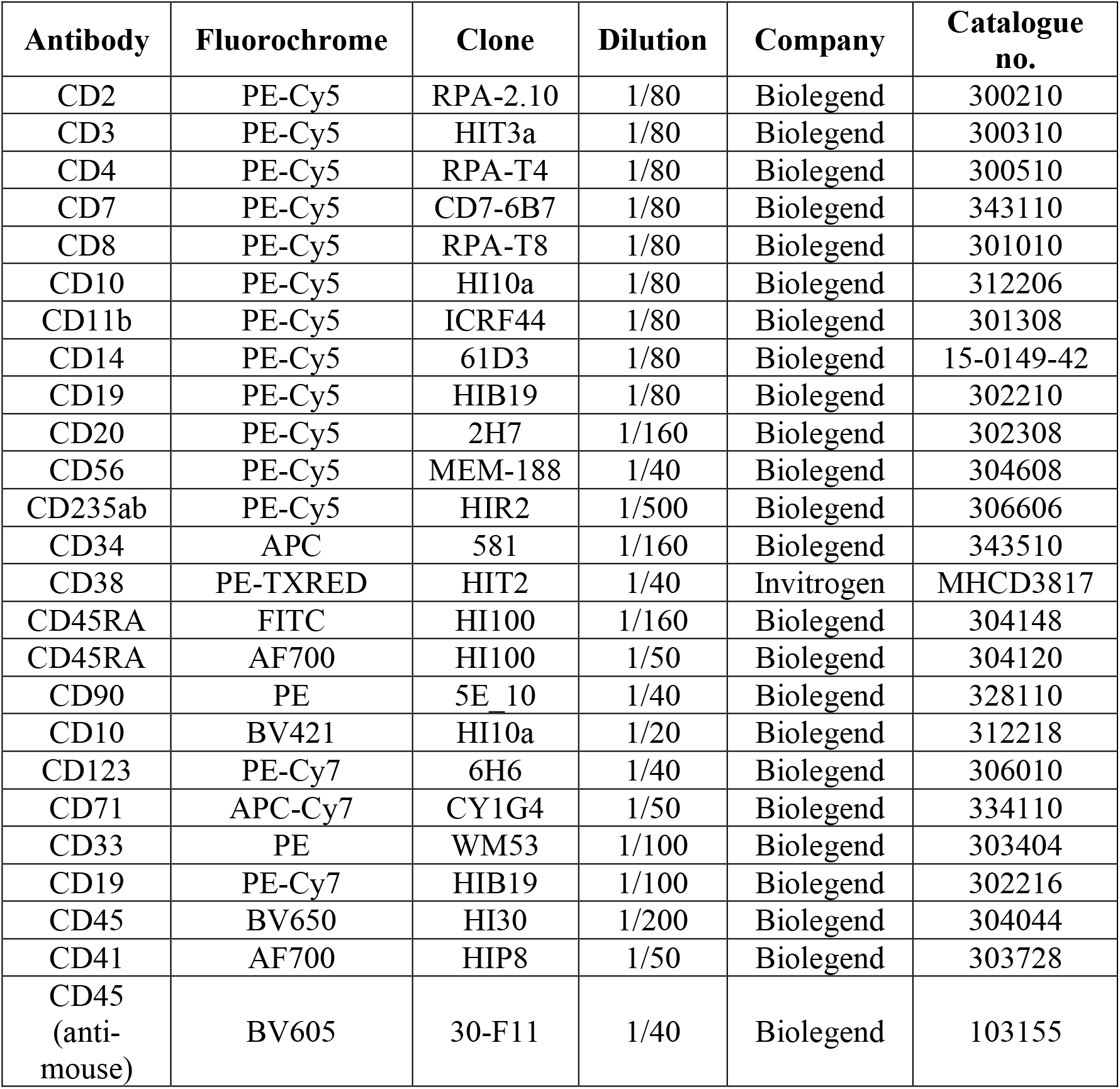
List of antibodies used for FACS staining.

**Table S3.**
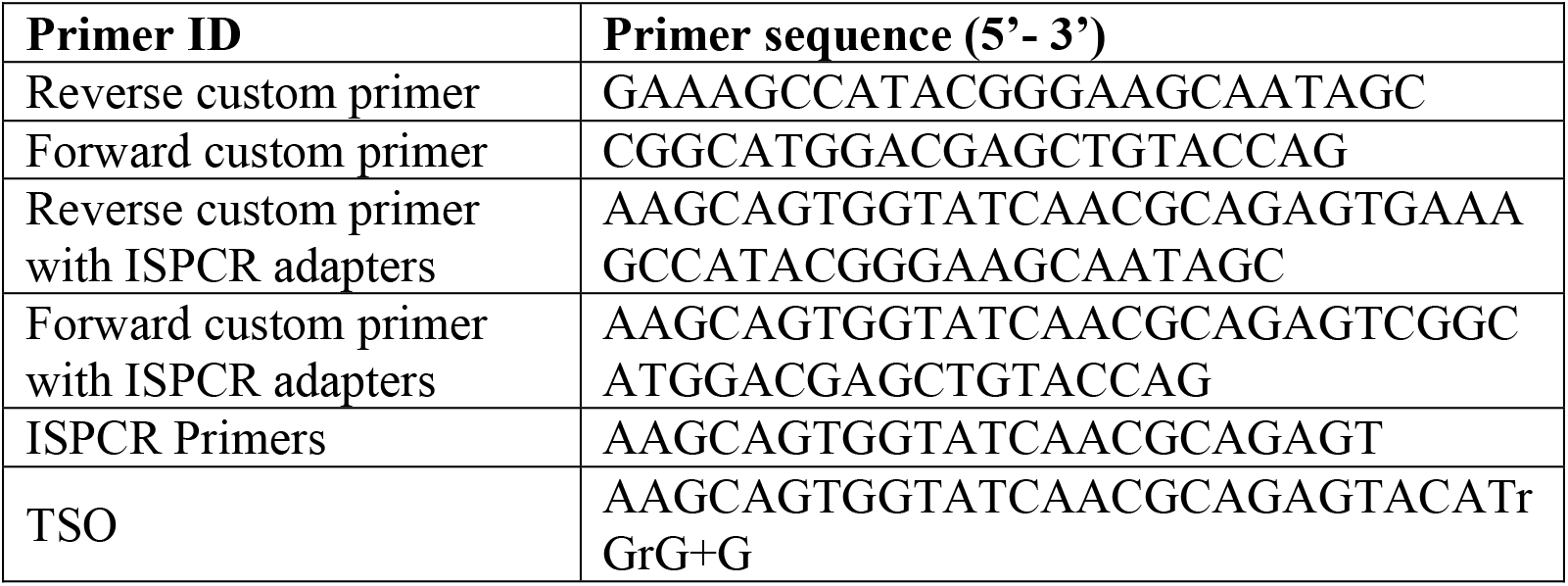
List of primers used in single-cell and bulk RNA sequencing.

## Supplemental datasets

**Data S1:** Genes differentially expressed between platelet-biased and multilineage hHSCs defined by clustering.

**Data S2:** Genes differentially expressed between platelet-biased and multilineage hHSCs defined by barcoding.

**Data S3:** Gene sets used for GSEA analysis

## References

1. S. E. W. Jacobsen, C. Nerlov, Haematopoiesis in the era of advanced single-cell technologies. Nat Cell Biol 21, 2–8 (2019).

2. B. Dykstra et al., Long-term propagation of distinct hematopoietic differentiation programs in vivo. Cell stem cell 1, 218–229 (2007).

3. R. Yamamoto et al., Clonal analysis unveils self-renewing lineage-restricted progenitors generated directly from hematopoietic stem cells. Cell 154, 1112–1126 (2013).

4. J. Carrelha et al., Hierarchically related lineage-restricted fates of multipotent haematopoietic stem cells. Nature 554, 106–111 (2018).

5. W. Pei et al., Polylox barcoding reveals haematopoietic stem cell fates realized in vivo. Nature 548, 456–460 (2017).

6. A. E. Rodriguez-Fraticelli et al., Clonal analysis of lineage fate in native haematopoiesis. Nature 553, 212–216 (2018).

7. A. E. Rodriguez-Fraticelli et al., Single-cell lineage tracing unveils a role for TCF15 in haematopoiesis. Nature 583, 585–589 (2020).

8. A. Grover et al., Single-cell RNA sequencing reveals molecular and functional platelet bias of aged haematopoietic stem cells. Nat Commun 7, 11075 (2016).

9. W. W. Pang et al., Human bone marrow hematopoietic stem cells are increased in frequency and myeloid-biased with age. Proc Natl Acad Sci U S A 108, 20012–20017 (2011).

10. S. Valletta et al., Micro-environmental sensing by bone marrow stroma identifies IL-6 and TGFbeta1 as regulators of hematopoietic ageing. Nat Commun 11, 4075 (2020).

11. T. N. Rao et al., JAK2-V617F and interferon-alpha induce megakaryocyte-biased stem cells characterized by decreased long-term functionality. Blood 137, 2139–2151 (2021).

12. J. Tong et al., Hematopoietic Stem Cell Heterogeneity Is Linked to the Initiation and Therapeutic Response of Myeloproliferative Neoplasms. Cell Stem Cell 28, 502–513 e506 (2021).

13. F. Notta et al., Isolation of single human hematopoietic stem cells capable of long-term multilineage engraftment. Science 333, 218–221 (2011).

14. E. Laurenti et al., CDK6 levels regulate quiescence exit in human hematopoietic stem cells. Cell Stem Cell 16, 302–313 (2015).

15. A. M. Cheung et al., Analysis of the clonal growth and differentiation dynamics of primitive barcoded human cord blood cells in NSG mice. Blood 122, 3129–3137 (2013).

16. M. E. Belderbos et al., Donor-to-Donor Heterogeneity in the Clonal Dynamics of Transplanted HumanCord Blood Stem Cellsin Murine Xenografts. Biol Blood Marrow Transplant 26, 16–25 (2020).

17. L. Biasco et al., In Vivo Tracking of Human Hematopoiesis Reveals Patterns of Clonal Dynamics during Early and Steady-State Reconstitution Phases. Cell Stem Cell 19, 107–119 (2016).

18. E. Six et al., Clonal tracking in gene therapy patients reveals a diversity of human hematopoietic differentiation programs. Blood 135, 1219–1231 (2020).

19. A. Rundberg Nilsson, S. Soneji, S. Adolfsson, D. Bryder, C. J. Pronk, Human and Murine Hematopoietic Stem Cell Aging Is Associated with Functional Impairments and Intrinsic Megakaryocytic/Erythroid Bias. PLoS One 11, e0158369 (2016).

20. M. E. Belderbos et al., Clonal selection and asymmetric distribution of human leukemia in murine xenografts revealed by cellular barcoding. Blood 129, 3210–3220 (2017).

21. B. E. McIntosh et al., Nonirradiated NOD,B6.SCID Il2rgamma-/-Kit(W41/W41) (NBSGW) mice support multilineage engraftment of human hematopoietic cells. Stem Cell Reports 4, 171–180 (2015).

22. S. Pinho et al., Lineage-Biased Hematopoietic Stem Cells Are Regulated by Distinct Niches. Dev Cell 44, 634–641 e634 (2018).

23. M. Zhao et al., Megakaryocytes maintain homeostatic quiescence and promote post-injury regeneration of hematopoietic stem cells. Nat Med 20, 1321–1326 (2014).

24. C. Di Genua et al., C/EBPalpha and GATA-2 Mutations Induce Bilineage Acute Erythroid Leukemia through Transformation of a Neomorphic Neutrophil-Erythroid Progenitor. Cancer Cell 37, 690–704 e698 (2020).

25. S. Picelli et al., Full-length RNA-seq from single cells using Smart-seq2. Nat Protoc 9, 171–181 (2014).

26. A. Rodriguez-Meira et al., Unravelling Intratumoral Heterogeneity through High-Sensitivity Single-Cell Mutational Analysis and Parallel RNA Sequencing. Mol Cell 73, 1292–1305 e1298 (2019).

27. A. Giustacchini et al., Single-cell transcriptomics uncovers distinct molecular signatures of stem cells in chronic myeloid leukemia. Nat Med 23, 692–702 (2017).

28. F. Krueger. (2012).

29. A. Dobin et al., STAR: ultrafast universal RNA-seq aligner. Bioinformatics 29, 15–21 (2013).

30. Y. Liao, G. K. Smyth, W. Shi, The Subread aligner: fast, accurate and scalable read mapping by seed-and-vote. Nucleic Acids Res 41, e108 (2013).

31. P. Ewels, M. Magnusson, S. Lundin, M. Kaller, MultiQC: summarize analysis results for multiple tools and samples in a single report. Bioinformatics 32, 3047–3048 (2016).

32. Y. Hao et al., Integrated analysis of multimodal single-cell data. Cell 184, 3573-3587.e3529 (2021).

33. R. B. Patterson-Cross, A. J. Levine, V. Menon, Selecting single cell clustering parameter values using subsampling-based robustness metrics. BMC Bioinformatics 22, 1–13 (2021).

34. S. Aibar et al., SCENIC: single-cell regulatory network inference and clustering. Nature Methods 14, 1083–1086 (2017).

35. A. Giladi et al., Single-cell characterization of haematopoietic progenitors and their trajectories in homeostasis and perturbed haematopoiesis. Nature Cell Biology 20, 836–846 (2018).

36. F. K. B. Lauridsen et al., Differences in Cell Cycle Status Underlie Transcriptional Heterogeneity in the HSC Compartment. Cell Reports 24, 766–780 (2018).

37. I. Tirosh et al., Dissecting the multicellular ecosystem of metastatic melanoma by single-cell RNA-seq. Science 352, 189–196 (2016).

38. N. K. Wilson et al., Combined Single-Cell Functional and Gene Expression Analysis Resolves Heterogeneity within Stem Cell Populations. Cell Stem Cell 16, 712–724 (2015).

39. E. M. Pietras et al., Functionally Distinct Subsets of Lineage-Biased Multipotent Progenitors Control Blood Production in Normal and Regenerative Conditions. Cell Stem Cell 17, 35–46 (2015).

40. S. Durinck, P. T. Spellman, E. Birney, W. Huber, Mapping identifiers for the integration of genomic datasets with the R/ Bioconductor package biomaRt. Nature Protocols 4, 1184–1191 (2009).

41. S. Belluschi et al., Myelo-lymphoid lineage restriction occurs in the human haematopoietic stem cell compartment before lymphoid-primed multipotent progenitors. Nat Commun 9, 4100 (2018).

42. F. K. B. Lauridsen et al., Differences in Cell Cycle Status Underlie Transcriptional Heterogeneity in the HSC Compartment. Cell Rep 24, 766–780 (2018).

43. A. Liberzon et al., The Molecular Signatures Database (MSigDB) hallmark gene set collection. Cell Syst 1, 417–425 (2015).

44. G. Korotkevich et al., Fast gene set enrichment analysis. bioRxiv, (2021).

45. S. I. Shen Li. (Bioconductor version: Release (3.14), 2021).

